# Novel Evoked Synaptic Activity Potentials (ESAPs) elicited by Spinal Cord Stimulation

**DOI:** 10.1101/2023.02.18.528981

**Authors:** Mahima Sharma, Vividha Bhaskar, Lillian Yang, Mohamad FallahRad, Nigel Gebodh, Tianhe Zhang, Rosana Esteller, John Martin, Marom Bikson

## Abstract

Spinal cord stimulation (SCS) evokes fast epidural Evoked Compound Action Potential (ECAPs) that represent activity of dorsal column axons, but not necessarily a spinal circuit response. Using a multimodal approach, we identified and characterized a delayed and slower potential evoked by SCS that reflects synaptic activity within the spinal cord. Anesthetized female Sprague Dawley rats were implanted with an epidural SCS lead, epidural motor cortex stimulation electrodes, an epidural spinal cord recoding lead, an intraspinal penetrating recording electrode array, and intramuscular electromyography (EMG) electrodes in the hindlimb and back. We stimulated the motor cortex or the epidural spinal cord and recorded epidural, intraspinal, and EMG responses. SCS pulses produced characteristic propagating ECAPs (composed of P1, N1, and P2 waves with latencies <2 ms) and an additional wave (“S1”) starting after the N2. We verified the S1-wave was not a stimulation artifact and was not a reflection of hindlimb/back EMG. The S1-wave has a distinct stimulation-intensity dose response and spatial profile compared to ECAPs. CNQX (a selective competitive antagonist of AMPA receptors) significantly diminished the S1-wave, but not ECAPs. Furthermore, cortical stimulation, which did not evoke ECAPs, produced epidurally detectable and CNQX-sensitive responses at the same spinal sites, confirming epidural recording of an evoked synaptic response. Finally, applying 50 Hz SCS resulted in dampening of ESAPs, but not ECAPs. Therefore, we hypothesize that the S1-wave is synaptic in origin, and we term the S1-wave type responses: Evoked Synaptic Activity Potentials (ESAPs). The identification and characterization of epidurally recorded ESAPs from the dorsal horn may elucidate SCS mechanisms.

**Significance Statement:** Spinal cord stimulation (SCS) is an established treatment for chronic pain and has applications to other disorders and neurorehabilitation. Notwithstanding decades of trials and research, questions remain about SCS mechanisms of action - and indicators thereof. Recent technological developments have enabled the detection of Evoked Compound Action Potential (ECAPs) – reflecting synchronous activity of the dorsal column axons activated by SCS. However, ECAP is not a direct measure of sensory processing in the dorsal horn. Here, we identify and characterize a novel electrophysiological signal that is evoked and detectable by epidural SCS electrodes and reflects spinal synaptic currents. This new signal, termed an Evoked Synaptic Activity Potential (ESAP), is thus a novel means with which to interrogate spinal gray matter circuits during SCS.

## Introduction

Epidural spinal cord stimulation (SCS) for neuropathic pain is thought to engage inhibitory dorsal horn mechanisms, whether via synaptic activation following antidromic stimulation of dorsal column fibers, consistent with Gate Control Theory (Shealy et al., 1970), or through more subtle, slower acting mechanisms (Titus et al., 2021). Dorsal column stimulation is sensed by evoked electrical responses representing the summed activity of dorsal column (Aβ) axons, whose synchronous activation produces a propagating evoked compound action potential (ECAP) characterized by P1, N1, and P2 waves, with latencies <2 ms proximal to the stimulation site (Cedeño et al., 2022; Cioni and Meglio, 1986; Dietz et al., 2022; Parker et al., 2013). The stimulation current threshold for ECAPs correlates with patient reported paresthesia and discomfort thresholds, and ECAP recording can be performed using contacts on the therapeutic stimulation leads, supporting the use of ECAPs in closed-loop SCS systems (Parker et al., 2012). However, dorsal column activity (i.e., the ECAP) is not itself directly indicative of pain or analgesia (Pilitsis et al., 2021; Vallejo et al., 2021), inviting a search for other signals that may better indicate pain relief by SCS.

Experimental electrophysiology involving the spinal cord, spanning decades, identified field potentials beyond the ECAP with slower time-courses ranging from a few ms to tens or hundreds of ms after a given stimulus (Cervero et al., 1978; Ondrejcák et al., 2005; Wall, 1958; Yates et al., 1982). Historically, studies of slow spinal potentials supported development of theories for pain manifestation and its control (Wall, 1958), including Gate Control Theory, and initial mechanistic hypotheses about SCS were tested through measurement of longer-latency “prolonged small fiber afterdischarge” (PSAD) (Shealy et al., 1970). Slow spinal potentials may correlate with modulation of afferent input (e.g. primary afferent depolarization), interneuron activity, or dorsal horn excitability (Contreras-Hernández et al., 2022, 2015; Manjarrez et al., 2003; Wall, 1958) and may therefore provide more information about the spinal (pain) state than ECAPs alone. However, fundamental studies on slower potentials used electrodes not suitable for clinical applications. Analysis of epidurally recorded evoked potentials for SCS has focused on the fast ECAPs (Anaya et al., 2020; Parker et al., 2012); slower evoked responses have been sparsely noted but unreliably, at high clinical intensities, and topically described as signs of patient discomfort (Parker et al., 2012) or muscle activation (Falowski et al., 2022).

Here we use a rodent model to identify and characterize an electrophysiological signal evoked and detectable by epidural SCS electrodes yet reflecting intra-spinal synaptic activity, and with a threshold above ECAP thresholds but below EMG thresholds, and a latency after ECAP onset but before EMG onset. We show this S1-wave reflects grey-matter excitatory synaptic currents, as part of a type of responses that we term Evoked Synaptic Activity Potentials (ESAPs). The distinct etiology of ESAPs (grey matter synaptic processing) vs. ECAPs (white matter conduction) is reflected in distinct (spatio-temporal, stimulation intensity, single pulse vs. tonic) responses to SCS. Building on the work described here, ongoing studies on whether and how ESAPs indicate effects of SCS and/or dorsal horn state (e.g., sensory/proprioceptive processing) are warranted.

## Methods

### Ethics Statement

All experiments were performed in accordance with the *NIH’s Guide for the Care and Use of Laboratory Animals*. Care and treatment of the animals conformed to protocols approved by the Institutional Animal Care and Use Committee of the City University of New York Advanced Science Research Center.

### Animal Subjects

A total of 47 adult female Sprague-Dawley rats, weighing 250-300 g on the day of the experiment, were used for all procedures. Rats were housed in groups of two to three per filter-top polycarbonate cages at our climate-controlled vivarium with the standard 12 h light –dark cycle, room temperature of 22.5 ± 0.9 °C, relative humidity of 36 ± 4 %, food and water *ad libitum*. Rats had a minimum 3-day acclimation period before starting any experiment. All procedures were conducted between 10:00 and 17:00 hours local time.

### Anesthesia and animal preparation

Rats were anesthetized using intraperitoneal (IP) injection of a mixture of ketamine and xylazine (70 mg/kg; 6 mg/kg). Care was taken to maintain a relatively stable anesthetic state and body temperature. Body temperature was monitored with a rectal probe and maintained at 37 ± 1°C during the entire experimental procedure, using a feedback controlled warming pad (Physio Suite, Kent Scientific). Ophthalmic ointment was applied to the eyes to prevent ocular dryness. The scalp, back and left hindlimb were shaved to facilitate craniotomy, laminectomy, and optimal measurements from EMG recording electrodes.

To assess the appropriate depth of anesthesia, periodic monitoring was conducted of respiratory and heart rates (Physio Suite, Kent Scientific), muscle tone, the absence of vibrissae whisking, and the absence of hindlimb withdrawal to foot pinch. The required anesthetic depth was maintained throughout the experiment with regular IP administration of supplemental doses of ketamine (25-35 mg/kg) every 90 minutes (Borrell et al., 2017).

#### Surgery

All the rats were subjected to laminectomy; 10 rats also underwent a craniotomy. During all surgical procedures, hemostasis was generally achieved through gentle pressure and the wounds were irrigated using sterile saline. Warm sterile saline was applied to surgical wounds to protect the tissues from desiccation during the entire experimental procedure.

##### Laminectomy

Rats were placed in a stereotaxic frame (David Kopf Instruments) after induction of anesthesia. Following shaving of the back and subcutaneous administration of 0.5% lidocaine (Henry Schein) at the incision site, a midline incision was made using the scalpel blade over the skin of the back to expose the T10-L2 vertebrae. The overlying musculature and fascia were cleared. After identifying the T13 vertebrae, a bilateral laminectomy was performed at the T10 and T11 vertebrae, to expose T12-L1 spinal segments. A hemi-laminectomy was performed on the left side of L1 and bilateral laminectomy at the L2 vertebrae to expose the L4-S3 segments of the spinal cord. Muscles on each side of midline were expanded slowly using a self-retaining retractor placed under the muscle. To stabilize the spinal cord for neural recordings, the L3 lamina was clamped. An incision was made in the dura at L4/L5 spinal segments and the pia was microdissected for insertion of the intraspinal electrode array and/or intrathecal drug administration.

##### Craniotomy

After placing the rats in a stereotaxic frame (David Kopf Instruments) and adjusting the incisor bar to ensure a flat skull position (equal heights of lambda and bregma skull points), a midline scalp incision was made. Craniotomy was performed over the hindlimb representation of the motor cortex (1.7 ± 0.4 mm posterior to bregma, 1.5 ± 0.5 mm lateral to midline) by thinning the skull over the primary motor cortex (5 mm X 5 mm) using a small burr. The thinned bone flap was then peeled off using fine micro-forceps. The general location of the craniotomy was guided by previous motor mapping studies of the rat (Fonoff et al., 2009; Frost et al., 2013) and was made intentionally larger than the anticipated location of the hindlimb motor cortex to account for slight variability among animals and ensure ample space for the electrode placement.

##### Placement of epidural spinal leads

We utilized two custom-made cylindrical 4-electrode leads (0.5 mm diameter; 1 mm inter-electrode distance; 0.5 mm wide; Boston Scientific Neuromodulation; Fig. 1C, 1D) – one to stimulate the spinal cord and other to record the epidural spinal responses. Both leads were placed epidurally over the midline of dorsal spinal cord. The epidural stimulation lead was inserted in the caudal direction through the laminectomy site at T10-T11 vertebral levels. The stimulation lead was adjusted so that active contacts were at the T12/T13 vertebral levels analogous to previous studies (Shechter et al., 2013; Tao et al., 2021). Vertebral locations were chosen based on prior work demonstrating that vertebral levels T12-L1 correspond to lumbar spinal dermatomes (L2-L5) often explored in chronic pain and motor systems research (Gelderd and Chopin, 1977; Gerasimenko et al., 2019; Swett and Woolf, 1985). The epidural recording lead was inserted in the rostral direction through the laminectomy site at L1 vertebra. The recording lead was adjusted so that the contacts were at T13/L1 vertebral levels to record dorsal epidural spinal responses from spinal lumbar segments L2-L5 (Dietz et al., 2022).

**Figure 1:**
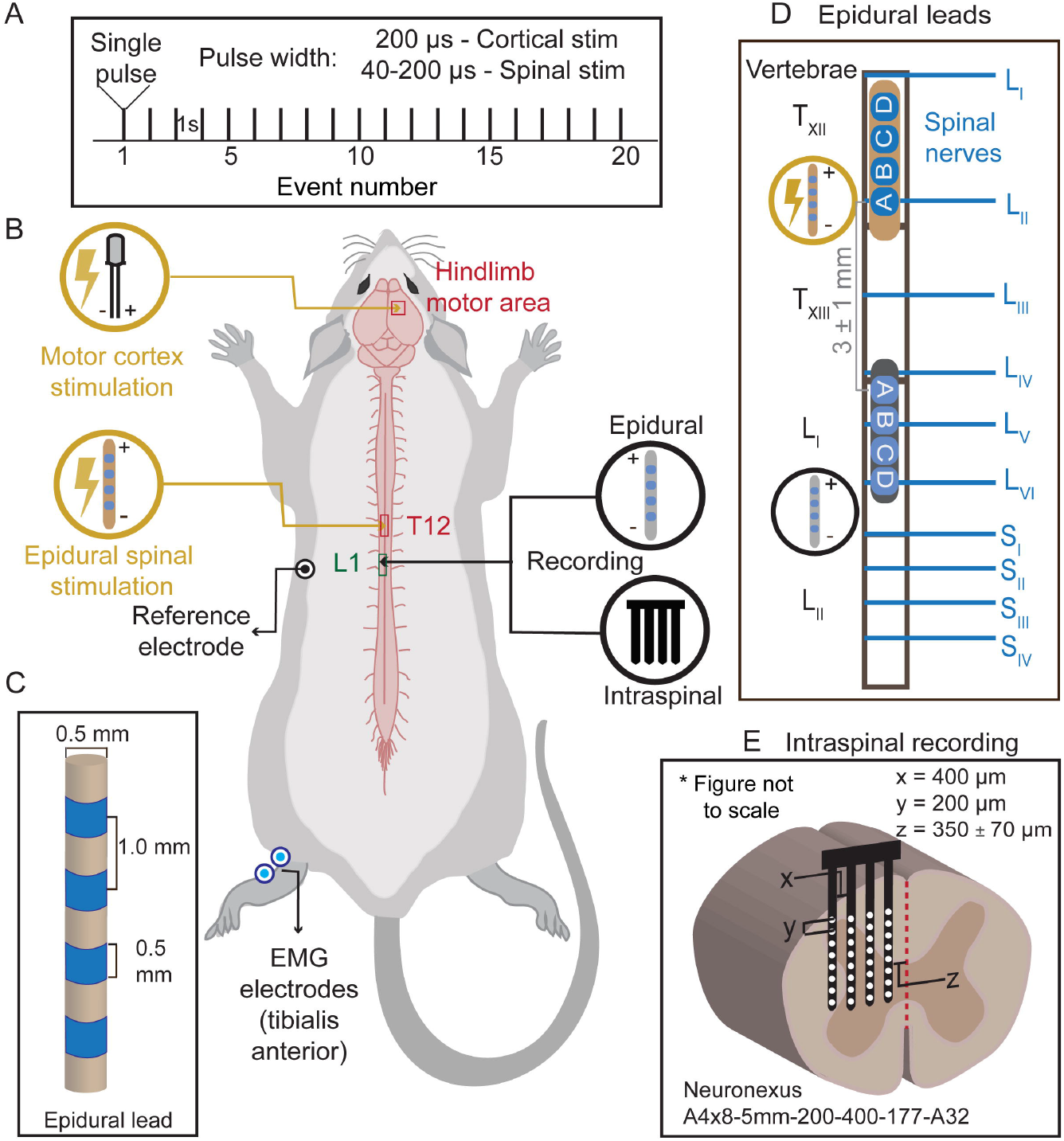
Experimental setup combining epidural cortical stimulation, epidural spinal stimulation, epidural spinal recording, invasive spinal recording, and EMG in rodent model. **(A)** Representative cortical and epidural spinal stimulation pulse trains of 20 low-frequency pulses, with pulse width of 200 µs per phase and/or 40-200 µs per phase, respectively. **(B)** Rat model depicting different electrodes sites. The right hindlimb representation area (HLA) of the motor cortex (M1) is stimulated using epidural bipolar stimulating electrode. Epidural leads are positioned on the midline of the spinal cord at T12-T13 segment and L1 vertebral segment to stimulate (orange) and record (gray) from the spinal cord, respectively. A four-shank silicon array (black) records the intraspinal responses. A reference electrode is placed under the skin. Percutaneous nickel-chrome wire electrodes (deinsulated 1 mm from the tip) bilaterally implanted into the *tibialis anterior* muscles record EMG responses. **(C)** Epidural lead with four electrodes of 0.5 mm length each, separated by 1 mm. The diameter of the lead is 0.5 mm. **(D)** Representative placement of the 4-electrode epidural leads (contacts A, B, C, and D oriented as shown) along the spinal cord. Stimulating (orange) and recording (gray) epidural leads are placed at the T12 vertebral segment and L1 vertebral segment, respectively, with the inter-electrode distance between electrode contact A of the epidural stimulating lead and electrode contact A of the epidural recording lead of 3 ± 1 mm. **(E)** Cross section of spinal cord showing the placement of electrode for intraspinal recording. A four-shank Neuronexus silicon array is inserted transversely into the left side of the spinal cord to a depth of 1.7 ± 0.2 mm from the pial surface. *Figure Contributions:* MS and NG designed the figure.

#### Electrophysiology

##### Cortical and epidural spinal stimulation

For stimulation of both the hindlimb representation area (HLA) and epidural dorsal columns, a constant current stimulator (A-M systems Model 2100) was set to deliver constant current symmetric biphasic stimulation either through the M1 cortical electrode or the epidural spinal lead, respectively. The M1 bipolar cortical stimulating electrode (PlasticsOne) was placed epidurally over HLA. The electrode was placed in the center of the hindlimb field on the motor cortex (1.8 ± 0.4 mm posterior to bregma, 1.6 ± 0.3 mm lateral to midline (Barth et al., 1990; Frost et al., 2013).

The epidural spinal stimulation lead had four electrode contacts designated ‘A’, ‘B’, ‘C’, ‘D’; with ‘A’ electrode nearest lead terminus. The inter-electrode distance between the ‘A’ electrodes of the epidural stimulating lead and the epidural recording lead was 3 ± 1 mm (Fig. 1D). Stimulation was applied using adjacent bipolar electrodes, delivering biphasic pulses with either leading anodic (rostral to caudal: − +) or cathodic (rostral to caudal: + −; unless otherwise indicated).

We optimized electrode placement to normalize the dose used. At the start of experiments, brief trains of biphasic pulses (333 Hz, contact A-B) were applied, as previously described for M1 mapping experiments (Song and Martin, 2017; Takemi et al., 2017); current amplitude was incremented (up to 0.8 mA) until either contractions or movement of the left hindlimbs were visually observed. The spinal stimulation lead was repositioned caudally/rostrally in ∼1 mm increments, as necessary, to elicit a specific visible movement of the hindlimb. Next, low-frequency epidural SCS was delivered through the designated stimulation lead while recording from the other lead (Fig. 1A). The stimulation or recording leads were repositioned caudally/rostrally in ∼1 mm increments and/or different electrode pairs were tested, as necessary, to determine the best site for stimulation and recording of a delayed response (e.g., S1 wave).

##### Epidural recording

The design of the epidural spinal lead used for recording was the same as the epidural lead for stimulation. The epidural recording lead was connected to a differential AC amplifier (A-M systems; model 1700) through customized connectors that allowed concurrent 100x amplification of three differential recordings (A-B, B-C, C-D; Fig. 2) from the four electrodes of the epidural lead. Signals were high pass filtered at 0.1 Hz and low pass filtered at 5 kHz. Signals were digitized (Cambridge Electronic Design 1401; RRID: SCR_017282), and recorded along with stimulation onset triggers (Cambridge Electronic Design Spike2 software; RRID: SCR_000903) at a sampling rate of 33333 Hz.

**Figure 2:**
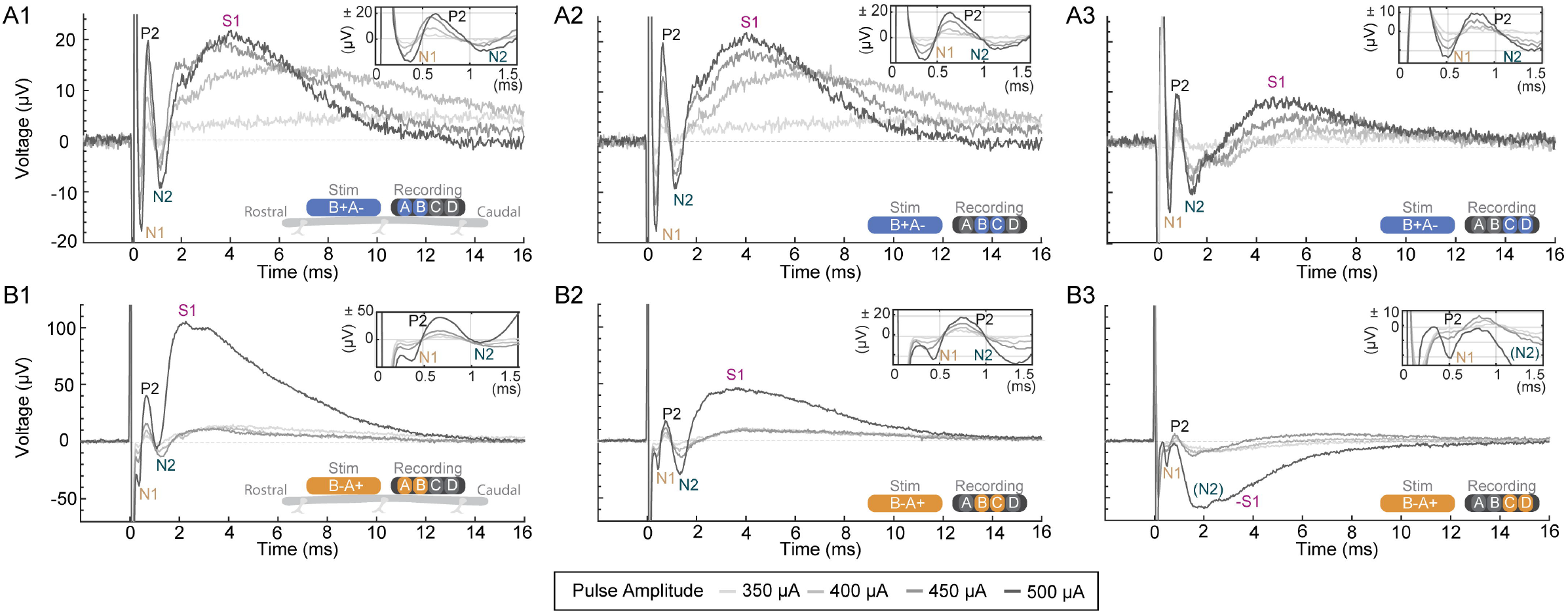
Representative stimulation amplitude vs. signal relationships (“dose responses”) of spinal potentials across epidural electrode contracts. The schematic at the lower right side of each panel shows the relative orientation and polarity of the stimulating electrodes (‘Stim’), for a proximal cathode (**A**, blue) or proximal anode (**B**, orange) stimulation polarity, as well as the recording electrodes (‘Recording’). Each panel shows the average evoked response from 20 low-frequency pulses (40 µs pulse width). The top right inset in each panels magnifies the spinal response from 0 ms to 1.5 ms. Stimulation intensity is reflecting in gray scale (350 µA, lightest gray; to 500 µA, darkest gray). Identified components of the response are labeled for 500 µA, when observable, as P1, N1, P2, N2, and S1. (**A1**) Dose response for cathode proximal stimulation with A vs B recording. (**A2**) Dose response for cathode proximal stimulation with B vs C recording. (**A2**) Dose response for cathode proximal stimulation with C vs D recording. (**B1**) Dose response for anode proximal stimulation with A vs B recording. (**B2**) Dose response for anode proximal stimulation with B vs C recording. (**B3**) Dose response for anode proximal stimulation with C vs D recording. *Figure Contributions:* MS and VB performed the experiments; MS and VB analyzed the data. Graphics contribution by NG.

##### Multichannel intraspinal recording

Extracellular, local field potentials (LFPs) were recorded from the L4/L5 spinal segment using a four-shank, 32-channel silicon probe (A4×8-5mm-200–400-177-A32; NeuroNexus, Fig. 1E) optimized for monopolar recording, as previously described (Amer et al., 2021; Song and Martin, 2017). The NeuroNexus probe was inserted transversely into the left side of the spinal cord (contralateral to the cortical stimulation electrode; Fig. 1A) to a depth of 1.7 ± 0.2 mm from the pial surface, as measured by the initial pial contact with the electrode array. Electrode depth was controlled using a 100 micron resolution electrode manipulator (David Kopf Instruments). The lateral-right shank of the probe was positioned 350 ± 70 µm from the midline, and the electrode array was lowered, following dura- and pia-puncture. Each shank was 400 µm apart, with eight recording sites, each with a site area of 177 µm^2^, located 200 µm apart (Fig. 1E). Electrode impedance ranged from 500 to 700 kΩ. Electrode positioning was carefully performed in all animals under a dissecting microscope to minimize surface dimpling and electrode position differences amongst animals.

A unity gain head-stage was connected via flexible cables (A32-OM32 connector; Neuronexus) to the spinal recording array. The broadband signals were amplified (1,000 X) and low-pass filtered (0.1–300 Hz) for LFP recordings (OmniPlex-D system; Plexon; RRID: SCR_014803). The LFPs were digitized at a sampling frequency of 1 kHz and offline 60-Hz notch-filtered (iirnotch, MATLAB 2021b, MathWorks).

##### EMG recording

EMG was recorded from left *tibialis anterior* (TA) muscles in the hindlimb or the left *longissimus thoracis* (LT) muscles in the back to assay motor evoked potentials (MEPs) and to determine the timing of evoked epidural responses in relation to the MEPs. To record the EMG, percutaneous nickel-chrome wire electrodes were prepared by deinsulating 1 mm of the tip on each wire and folding the wire back on itself from the exposed tip (‘hook electrode’). The wires were inserted into the belly of *tibialis anterior* or *longissimus thoracis* muscles using a 30-gauge hypodermic needle and were positioned approximately 2 mm apart from each other. Electrode location was verified by delivering a series of biphasic pulses to the dorsal epidural SCS electrode observing an EMG response through the inserted EMG electrodes.

For hindlimb EMG experiments, the presence of MEPs were monitored in response to the train of pulses to the contralateral M1 HLA or dorsal spinal cord. For back EMG experiments, the presence of MEPs were monitored in response to the train of pulses through the epidural electrodes. EMG data was continuously acquired with a differential AC amplifier system (A-M Systems, Model 1700), amplified at a gain of 1000, high pass filtered at 300 Hz and low pass filtered using second order Butterworth filter at 5 kHz (hindlimb EMG from TA) or 20 kHz (back EMG from LT). Signals were digitized using a CED 1401 data acquisition system (Cambridge Electronic Design), and recorded using Spike2 software (Cambridge Electronic Design) at a sampling rate of 33333 Hz, and rectified. EMG threshold is defined as the current amplitude that evoked responses with an amplitude greater than three times the standard deviation (SD) of the background signal level in 20% of the trials. EMG amplitudes are reported as root mean square (RMS) over 1-20 ms for initial hindlimb EMG, 25-45 ms for late hindlimb EMG and 1-9.9 ms for back EMG.

#### Drug Administration

To confirm the synaptic nature of the delayed wave component of the spinal potential, a competitive AMPA receptor antagonist, 6-cyano-7-nitroquinoxaline-2,3-dione (CNQX, 100 µl, 100 µM to 1 mM) was applied to the exposed spinal cord where the dura and pia were punctured at the L4/L5 spinal segment. 10 mM stock solutions of CNQX disodium salt hydrate (Sigma-Aldrich, Cat No. C239) were prepared by dissolving in 30 % DMSO solution. Stocks were then diluted in warm saline (37 ºC) to get the desired concentration of CNQX. The final concentration of DMSO in the solution never exceeded 0.1%. To control for the vehicle used in the experiments, warm saline solution (100 µl) was applied to the same site 20 minutes prior to the CNQX application. The vehicle-only experiment was done once or twice prior to the CNQX application, and sequential administration of CNQX enabled animals to be used as their own controls. The experiments with consistent evoked responses between the two saline conditions or the ones with no more than 40% change in the evoked response with saline application, were included for further analyses. After ∼20 minutes of drug application, warm saline (200 µl) was applied to wash out the CNQX. To assess the relationship between EMG and ESAP response further, the local anesthetic lidocaine (Henry Schein, 0.5%, 0.1 ml) was injected into the right and left LT muscles.

#### Data analysis

All signal processing in this study was performed with MATLAB (2021b, MathWorks; RRID: SCR_001622). Raw data was exported from Spike2 to MATLAB for analysis and following the offset correction, the average responses from the 20 trials or 40 trials (subset of animals where back EMG was recorded) were computed for each experiment. To characterize and compute the ECAP amplitude and the delayed wave, the positive peaks (P2, S1; Figures 2, 3, 5, 6) and the negative troughs (N1, N2) were located, and their amplitude and latencies were computed. The ECAP amplitude was always calculated as the peak-to-peak amplitude (voltage) between the first negative phase (N1) and the second positive peak (P2) of the ECAP. Latency was measured as the time from the onset of stimulus artifact to the onset of the first deflection of the potential.

**Figure 3:**
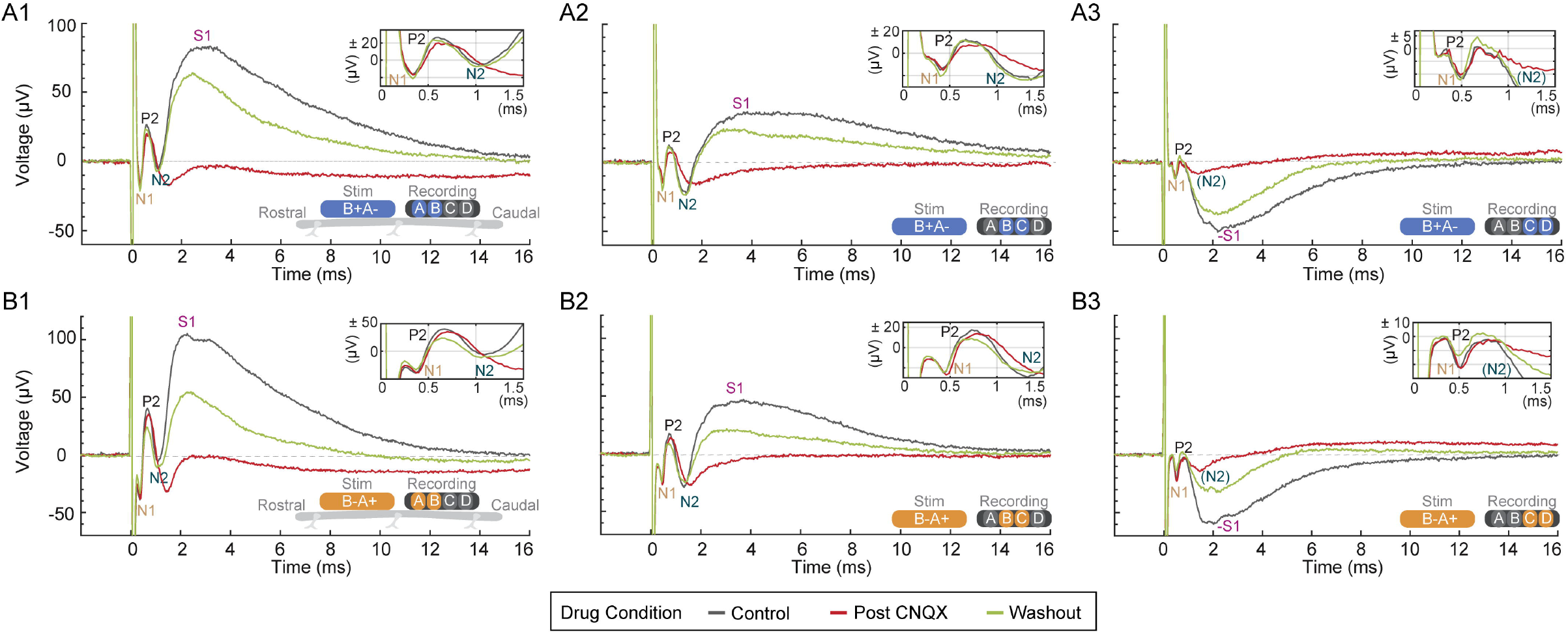
Dependence of evoked epidural spinal waveform on glutamatergic synapses. CNQX, a competitive AMPAR antagonist (1 mM, 100 µl) was applied to the superficial spinal cord surface. Evoked response are recoded before (control, gray), during (post CNQX, red) and after washout (green) of CNQX. The schematic at the lower right side of each panel shows the relative orientation and polarity of the stimulating electrodes (‘Stim’), for a proximal cathode (**A**, blue) or proximal anode (**B**, orange) stimulation polarity, as well as the recording electrodes (‘Recording’). Each panel shows the average evoked response from 20 pulses (40 µs pulse width). The top right inset in each panels magnifies the spinal response from 0 ms to 1.5 ms. Stimulation intensity was fixed across all panels. Identified waveform components of the response are labeled for the control condition, when observable, as P1, N1, P2, N2, and S1. (**A1**) CNQX effects for cathode proximal stimulation with A vs B recording. (**A2**) CNQX effects for cathode proximal stimulation with B vs C recording. (**A2**) CNQX effects for cathode proximal stimulation with C vs D recording. (**B1**) CNQX effects for anode proximal stimulation with A vs B recording. (**B2**) CNQX effects for anode proximal stimulation with B vs C recording. (**B3**) CNQX effects for anode proximal stimulation with C vs D recording. *Figure Contributions:* MS and VB performed the experiments; MS and VB analyzed the data.

EMG responses were rectified and root mean square (rms) and latency calculated. Data for different parameters of ECAP components, delayed wave and EMG are reported as mean ± SEM. Threshold for ECAP is defined as the minimum current amplitude that evoked responses in 95% of the trials that exceeded 10 µV. Threshold for the delayed wave component (S1) is defined as the minimum current amplitude that evoked responses in 95% of the trials that exceeded 25 µV.

##### Statistical analysis

Data were tested for normality using one sample Kolmogorov Smirnov test (kstest, MATLAB) or Shapiro-Wilk test (swtest, MATLAB). Statistical comparisons included parametric tests (ttest, MATLAB) or non-parametric tests (Wilcoxon signed rank test, signrank, MATLAB) for data sets with normal or non-normal distribution, respectively. One-way ANOVA (anova1, MATLAB) was used to compare the percent change in amplitudes and latencies of ECAP and S1 across low-frequency or tonic (50 Hz) SCS. A Bonferroni’s *post hoc* correction was used for multiple comparisons. Error bars indicate mean ± SEM. P values < 0.05 were considered significant. Further details on the comparisons made and statistical tests used are presented in Table 1. Custom MATLAB code that was used for analyses available upon request.

## Results

### Characterization of ECAP and S1 SCS Evoked Responses

We systematically explored the waveform of potentials evoked by a rostrally-positioned spinal epidural lead and recorded by a caudally-positioned spinal epidural lead (n=39). Stimulation was delivered in trains of single biphasic pulses (Fig. 1A) through varied stimulation or recording electrodes, and stimulation polarities and intensities (Fig. 1). Representative responses from one animal are shown (Fig. 2). Stimulation could evoke a characteristic multi-phasic early response, reported in human and other rodent studies, including a P1, N1, P2, and N2, which together form the ECAP, at a mean threshold of 237 ± 15 µA (n=39). The earliest component of the ECAP, the positive P1, was in some cases obscured by the stimulus artifact and was easiest to resolve with increasing inter-electrode distance (e.g., recording from electrode position C vs D) and with proximal anode stimulation polarity (Fig. 2 B3). The average P1 latency was 0.32 ± 0.02 ms (n=39). The next component was the negative N1 (0.56 ± 0.02 ms, n=39), followed by the positive P2 (0.87 ± 0.03 ms, n=39), and negative N2 (1.35 ± 0.04 ms, n=39). The relative latency of each of these components increased with distance from stimulation to recording electrode (Fig. 2, left to right columns) and when the distance from the proximal cathode was changed by reversing stimulation polarity, consistent with axonal propagation (Fig. 2, A vs B row). However, the polarity of N1, P2, and N2 did not invert with the stimulation artifact and was consistent across recording electrode pairs. In a minority of experiments, N2 was composed of dual peaks (not shown). Taken together, these recordings indicate that epidural stimulation was functional and could evoke ECAP responses indicative of dorsal column activity consistent with prior observations (Cedeño et al., 2022; Dietz et al., 2022).

We further identified a “S1” wave with a latency longer than N2 and a mean threshold of 307 ± 16 µA (n=39). Both ECAP and ESAP amplitude increased monotonically with increasing current intensities, but the ratio of ESAP to ECAP neither increased monotonically nor stayed at the same value across all stimulation amplitudes (Extended Data Fig. 1-1E), supporting a distinct origin for the ESAP. Electrode position and stimulation dose (intensity, polarity) response testing also identified distinct features of the S1-wave versus the other waveform components. The S1-wave appeared after N2 (2.99 ± 0.15 ms, n=39), although the initiation of the S1-wave appeared to overlap with the N2. In addition to starting later than the ECAP components, the S1-wave has a longer duration (6.34 ± 0.04 ms) as compared to ECAP (1.01 ± 0.02 ms, signed-rank P<0.0001, n=39), with a distinctive peak at higher current intensities (Example Fig. 6). Furthermore, unlike the ECAP, which retained a relatively consistent amplitude and morphology across differential pairs, the relative amplitude of S1-wave and ECAP changed with stimulation to recording electrode distance (Fig. 2, A1 vs A2 vs A3, B1 vs B2 columns) and could invert at caudal electrode sites (Fig. 2 B3). In such inversions, the polarity of the S1-wave was the same as N2. Inverting stimulation polarity altered S1 morphology on a given recording pair but did not necessarily invert the S1-wave, suggesting spatial dependence of S1 on the location of the stimulating cathode but ruling against S1 being a stimulation artifact (Fig. 2, A vs. B). Taken together, these results demonstrate a unique stimulation-response relationship and spatio-temporal profile for S1-waves compared to ECAPs, consistent with a distinct physiological source.

### Synaptic Origin of S1-waves (ESAPs)

The dependence of spinal evoked potentials on glutamatergic synaptic transmission was assessed using CNQX (Fig. 3), a competitive AMPAR antagonist (1 mM, 100 µl). Subdural superfusion of CNQX resulted in a significant attenuation of the S1-wave from 52.2 ± 19.1 µV (control, vehicle) to 22.8 ± 10.1 µV (15 min post CNQX, n=14) but did not affect the mean ECAPs (P2-N1: 57.5 ± 15.5 µV, control to 54.8 ± 15.7 µV, 15 min post CNQX). The mean S1-wave amplitude significantly decreased following CNQX (mean decrease of 62.2 ± 6.7 %; signed-rank P=0.0001, n=14), whereas the ECAP did not significantly decrease (10.5 ± 5.6 %; signed-rank P=0.0562, n=14). Washout with saline (vehicle) resulted in the recovery of the S1-wave amplitude after 45 minutes (Fig. 3), consistent with the reversible nature of the selective AMPA receptor blockade by CNQX. Given the distinct waveform following CNQX but *not* vehicle alone and strong selectivity for AMPA receptors by CNQX, we conclude that the S1 wave is a type of evoked spinal synaptic response, or “ESAP”. The origin and physiology of ESAPs are analyzed further in the next sections.

### Intraspinal analog of ESAPS

Simultaneous recording of epidural and intraspinal potentials during epidural spinal stimulation (Fig. 4, column B) showed an intraspinal analog of the S1-wave (ESAP) as well as the ECAP (called afferent terminal potentials; ATPS (Contreras-Hernández et al., 2022)). The mean latencies of the S1 peak and its intraspinal analog were 3.11 ± 0.16 ms and 2.78 ± 0.23 ms, respectively (t_(9)_ = 1.35, P=0.38, n=10). The intraspinal response was spatially specific and could reverse in polarity-consistent with intraspinal sinks/source. The intraspinal analog of the S1-wave was also suppressed by CNQX (Fig. 4 B3), further supporting the intraspinal synaptic origin of ESAPs. A later slow wave potential was also observed in both epidural (10-30 ms) and intraspinal recordings (15-30 ms). Taken together, these intraspinal recordings suggest local sites of sinks/sources underlying the S1 wave, consistent with spinal synaptic currents.

**Figure 4:**
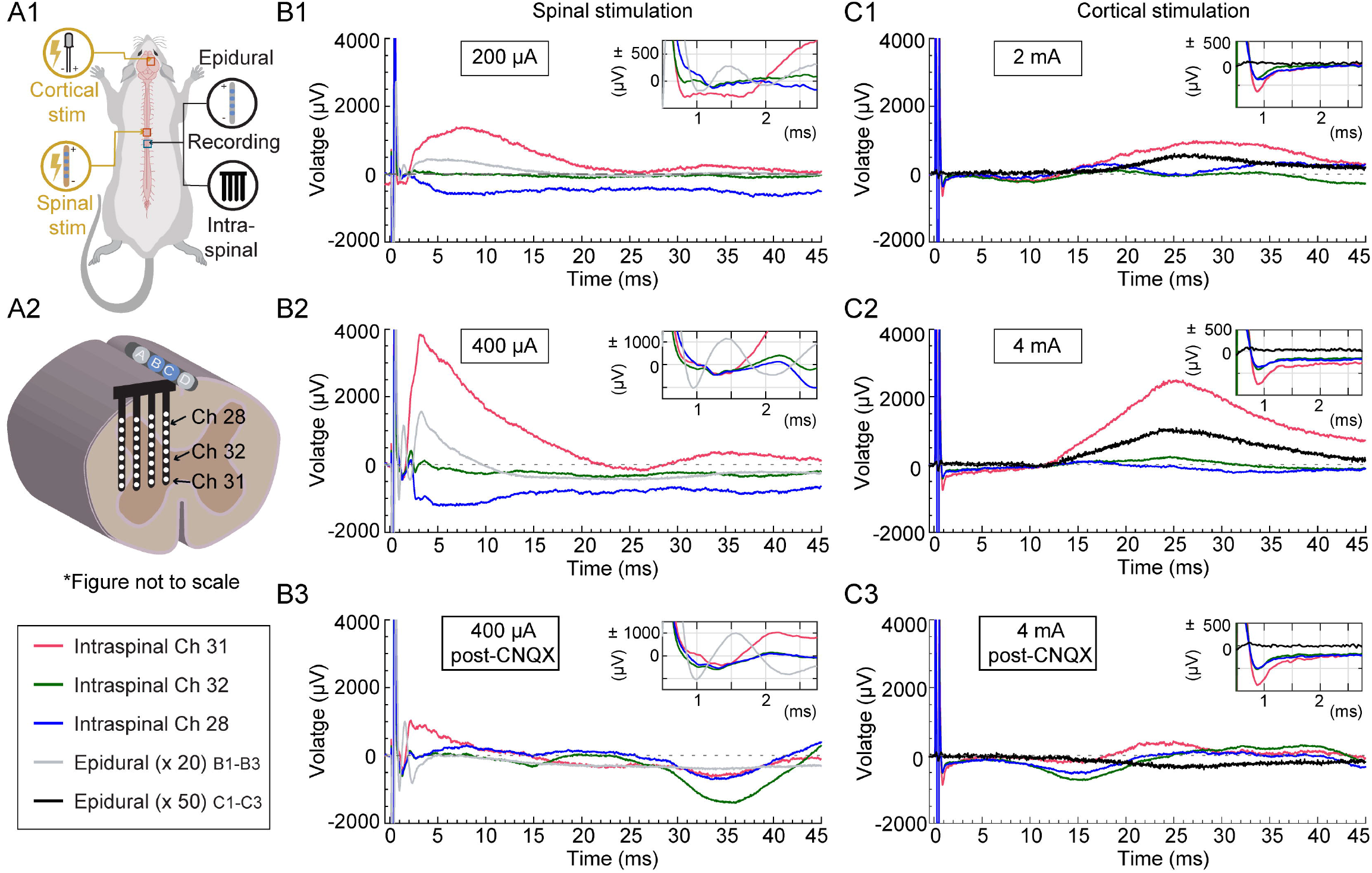
Simultaneous intraspinal and epidural recordings evoked with cortical and epidural spinal stimulation from the same rat. (A1) Rat model depicting location of different electrodes (both stimulating and recording, with 4 electrodes in total, Fig. 1D). (A2) Cross section of spinal cord with intraspinal electrode inserted transversely into the left side at a distance of 350 ± 70 µm from the epidural recording lead placed over the dorsal column with B and C (blue) as active recording electrodes. Intraspinal electrode array allows for recording from different sites: dorsal sector (Channel 28, blue), intermediate sector (Channel 32, black) (intermediate sector), and upper ventral sector (Channel 31, red) (upper ventral sector). Each panel shows the average evoked response from stimulation with 20 low-frequency pulses (200 µs pulse width) during different current intensities and conditions: **(B)** Epidural spinal stimulation at B1) 200 µA, B2) 400 µA, B3) 400 µA, 15 min post CNQX. Epidural recordings (gray) are magnified 20 times for better comparison. The top right insets in all panels show the spinal response from 0.5 ms to 2.7 ms post initiation of stimulation artifact. **(C)** Epidural cortical stimulation at C1) 2 mA, C2) 4 mA, C3) 4 mA, 15 min post CNQX. Epidural recordings (black) are magnified 50 times for better comparison. The top right insets in all panels show the spinal response from 0.5 ms to 2.7 ms post initiation of stimulation artifact. *Figure Contributions:* MS and LY performed the experiments; MS and NG analyzed the data.

We further considered the nature of the S1-waves and their intraspinal analog by evoking responses by epidural motor cortex stimulation. The spinal response to motor cortex stimulation reflects intra-spinal synaptic excitatory potentials (Amer et al., 2021; Zinger et al., 2013). Motor cortex stimulation produced a response recorded using epidural electrodes that was delayed and prolonged compared to spinal-stimulation evoked S1-waves (Fig. 4, column C), with a mean latency of 23.33 ± 0.90 ms. Intraspinal recordings reflect a similar waveform and latency of 22.53 ± 0.85 ms (t_(9)_ = 1.73, P=0.15, n=10), with spatial sensitivity (consistent with local intra-spinal sinks/sources). Cortical evoked responses were abolished by superfusion with CNQX (Fig. 4 C3). These results using cortical stimulation confirm intra-spinal (excitatory) signals can be detected with bipolar epidural cylindrical electrodes. The general correspondence between spinal responses to cortical vs. spinal stimulation should not imply the same synapses are active (only that they may overlap in location and type).

### Ruling out EMG as an Explanation for ESAP (S1)

The SCS current threshold, latencies, and behavior of the S1 ESAPs were compared to hindlimb/back EMG responses to assess whether or not S1 was directly related to EMG. In the subset of 9 animals from which both epidural and hindlimb EMG responses were recorded, mean S1 thresholds were significantly lower than hindlimb EMG thresholds (S1: 328 ± 28 µA vs. EMG: 694 ± 72 µA, ratio: 0.49 ± 0.04; t_(8)_ = 6.04, P<0.0001, n = 9). The current threshold for eliciting a visually-observed motor response was approximately double the hindlimb EMG threshold. As noted above, both ECAP and ESAP amplitude increased monotonically with stimulation current intensity (Fig. 5B1), but not ECAP to ESAP ratio (Extended Data Fig. 1-1C).

**Figure 5:**
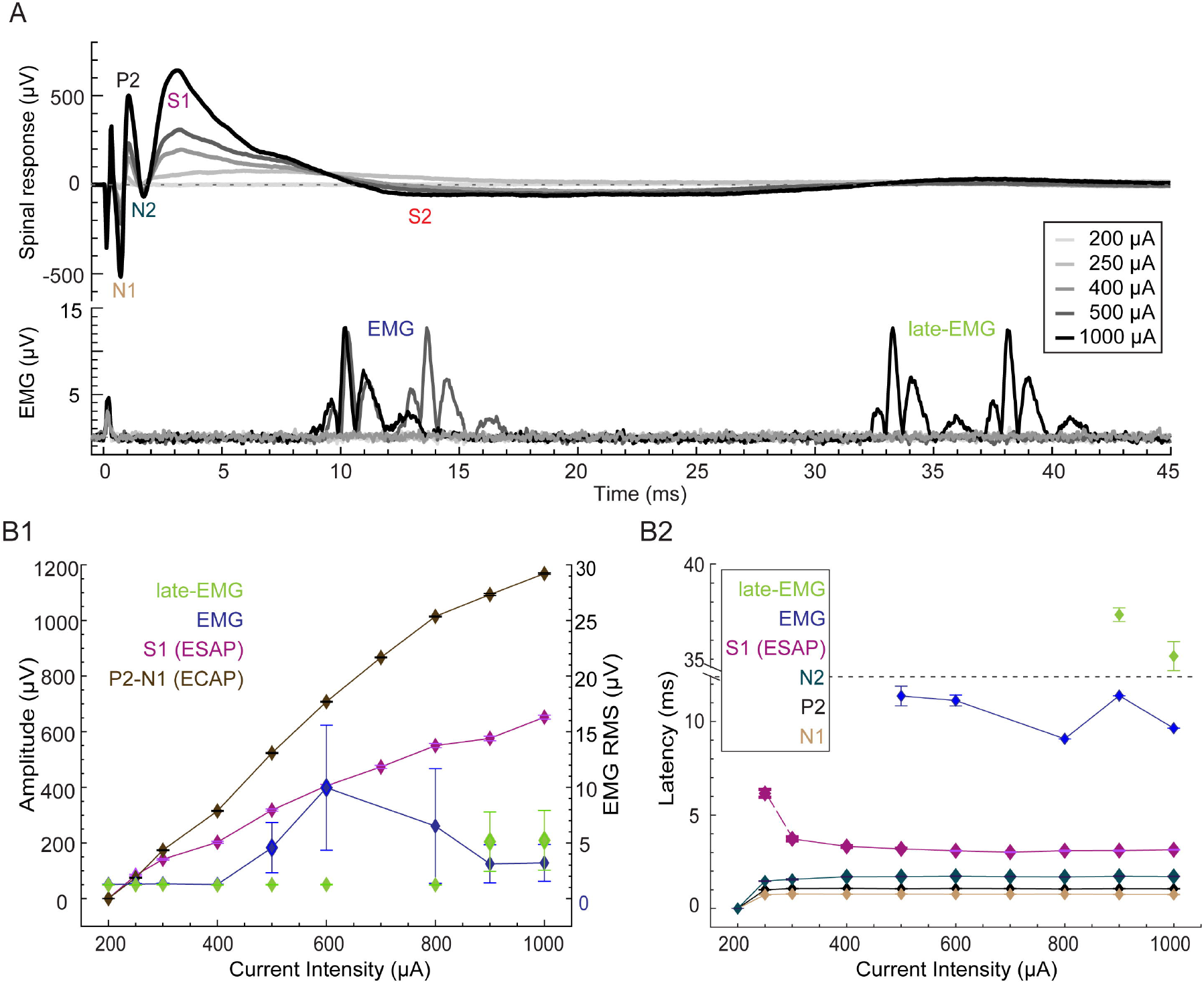
Dose response for the spinal and the hindlimb EMG recordings. **(A)** Upper panel shows the average evoked spinal response (ECAP, evoked synaptic activity potential ESAP (S1, S2)) and lower panel shows the average evoked EMG (*tibialis anterior* hindlimb muscle) response from 20 low-frequency pulses of epidural stimulation (Fig. 1A) of spinal cord at varying current intensities, ranging from 200 µA to 1000 µA) in a representative single rat. Identified waveform components of the spinal response are labeled for 1000 µA, when observable, as P1, N1, P2, N2, S1 & S2 (ESAP). ESAP (S1) becomes more distinct with increasing current intensity. Two types of EMG responses were observed: those occurring before 20 ms, referred to as EMG (blue), and those occurring after 30 ms, referred to as late-EMG (green). **(B1)** Dose-response curve for the ECAP (brown), ESAP (S1) (pink), EMG (blue) and late-EMG (green) responses. The left axis represents the amplitude for ECAP and ESAP (S1); the right axis represents the RMS for EMG responses. Each data point represents mean ± SEM (20 trials). **(B2)** Latencies plotted as a function of current intensities. N1 (brown), P2 (black), N2 (dark green), and S1 (ESAP, pink). Latency for the EMG and late-EMG is defined as the latency of the first peak in the EMG/late-EMG response. Data represented as mean ± SEM. *Figure Contributions:* MS and VB performed the experiments; MS analyzed the data.

With respect to latency, two distinct hindlimb EMG responses were evoked: A first EMG (∼9 ms) and a “late-EMG” (∼35 ms). Hindlimb EMG responses were stimulation-amplitude dependent. Across animals, ESAPs emerged gradually with increasing SCS amplitude (Extended Data Fig. 1-1B) and significantly earlier than first EMG (ESAP=3.06 ± 0.12 ms; EMG=9.37 ± 0.43 ms, t_(8)_ = 15.7, P<0.0001, n=9) and did not overlap in time with either the first or late EMG. The mean difference between the latencies for ESAP, measured at the S1 peak, and first hindlimb EMG, measured at the first rising edge of the signal, was 6.31 ± 0.4 ms (n=9).

In a separate experiment series, both epidural responses and back EMG (LT muscles) were recorded from a subset of 6 animals (Figure 6). No EMG was observed in 1 animal at up to the highest current amplitude tested (2000 µA). The mean S1 thresholds were significantly lower than back EMG thresholds (S1: 360 ± 40 µA vs. EMG: 1320 ± 239 µA, ratio: 0.32 ± 0.09 t_(4)_ = 4.19, P=0.01, n = 5). The current threshold for eliciting a visually-observed motor response was a further ∼1.7x the back EMG threshold. The mean difference between the latencies for ESAP, measured at the S1 peak (2.37 ± 0.16 ms), and the initiation of the (5.42 ± 0.8 ms), was 3.05 ± 0.8 ms (t_(2)_ = 3.7, P=0.06, n=3) (Figure 6A2). After injecting the local anesthetic lidocaine into the left and right LT muscle, EMG responses decreased 80 ± 1 % from 18.5 ± 6 µV RMS (pre-lidocaine) to 3.6 ± 1 µV RMS (20 min post lidocaine) (Figure 6B). There was no manifest change in the S1 peak amplitude (n=3); 269 ± 97 µV (pre-lidocaine), 204 ± 84 µV (20 min post lidocaine). In summary, the higher thresholds and longer latencies of EMG responses compared to ESAPs, as well as distinct waveforms and response to local anesthetic injection, further corroborate that ESAPs are not measuring muscle responses (i.e., ESAPS are not myogenic).

**Figure 6:**
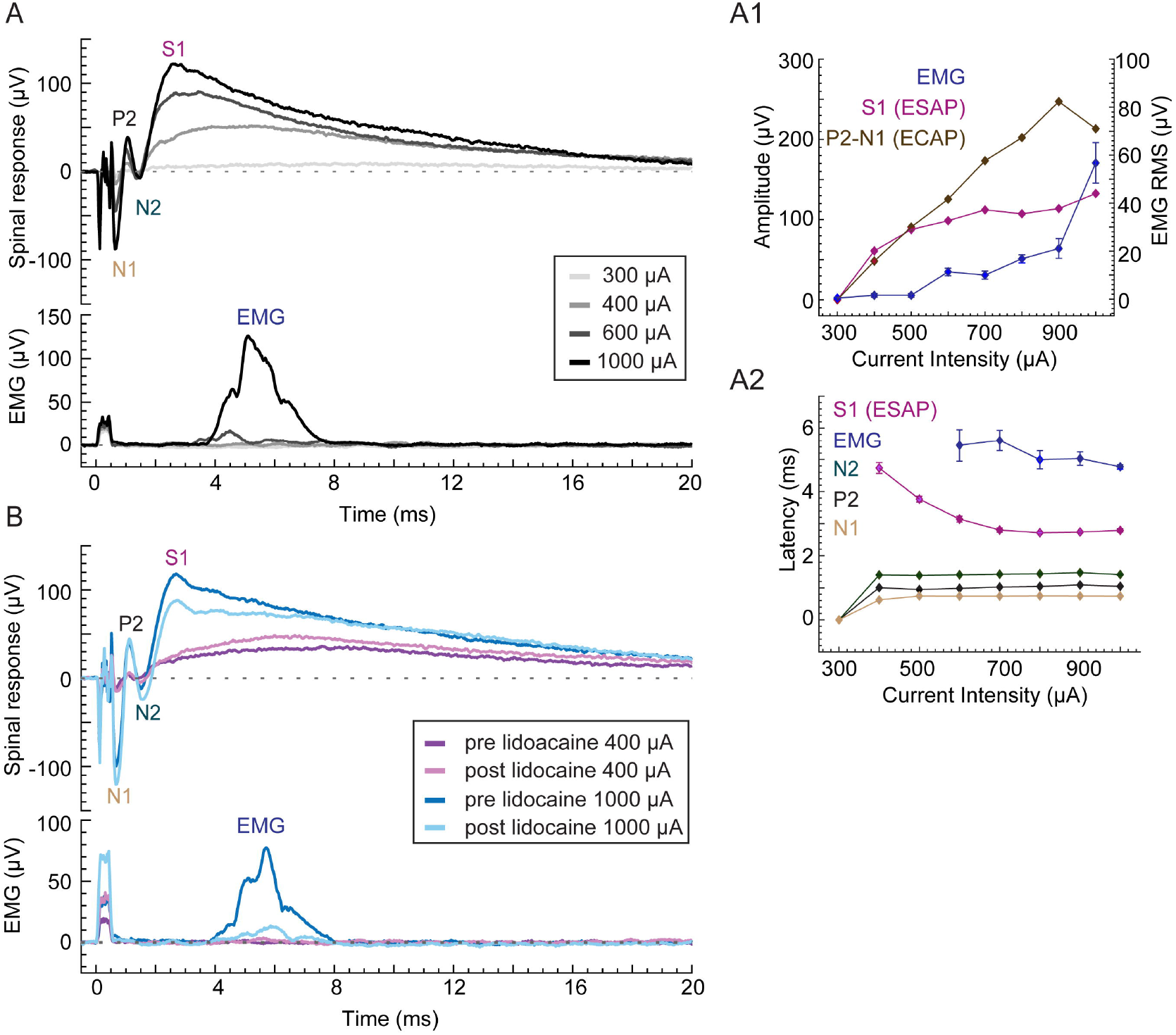
Dose response for the spinal and the back EMG recordings. **(A)** Upper panel shows the average evoked spinal response (ECAP, evoked synaptic activity potential ESAP (S1)) and lower panel shows the average evoked back EMG (LT muscle) response from epidural stimulation of spinal cord at varying current intensities, ranging from 200 µA to 1000 µA, with a train of 40 pulses (40 µs pulse width, 1 Hz) in a representative single rat. Identified waveform components of the spinal response are labeled for 1000 µA, when observable, as P1, N1, P2, N2, S1 & S2 (ESAP). ESAP (S1) becomes more distinct with increasing current intensity. **(A1)** Dose-response curve for the ECAP (brown), ESAP (S1) (pink), EMG (blue) and late-EMG (green) responses. The left axis represents the amplitude for ECAP and ESAP (S1); the right axis represents the RMS for EMG responses. Each data point represents mean ± SEM (40 trials). **(A2)** Latencies plotted as a function of current intensities. N1 (brown), P2 (black), N2 (dark green), and S1 (ESAP, pink). Latency for the back EMG is defined as the latency of the first peak in the EMG response. Data represented as mean ± SEM (40 trials). **(B)** Same as A, but only at two SCS current intensities-400 µA and 1000 µA, pre-lidocaine and 20 min post lidocaine in the same rat. *Figure Contributions:* MS performed the experiments and analyzed the data.

### Effects of SCS Frequency on ECAPs vs ESAPs (S1)

We evaluated the dependence of ECAPs and S1 ESAPs on the frequency of spinal cord stimulation (SCS) by evoking responses at 50 Hz SCS (Fig. 7). Responses are shown as a function of stimulation pulse index (event) to allow for comparison with evoked responses at low-frequency (1 Hz) SCS. One-way ANOVA revealed a significant decrease in the S1 amplitude (Fig. 7B, 7C) with increasing stimulation pulse count during 50 Hz SCS (F_(19,80)_=6.5162, P<0.001), unlike during 1 Hz SCS (F_(19,80)_=1.3284, P=0.1898; Fig. 8A). *Post-hoc* Bonferroni’s multiple comparison test demonstrated a significant change in S1 amplitude during 50 Hz SCS starting from event 10 (P=0.0082), with an average 51% decrease by event 14 (P=0.0007, n=5). In contrast, despite a non-significant trend toward facilitation in the first 9 events during 50 Hz SCS, there was no consistent significant change in mean ECAP amplitudes across stimuli events (Fig. 8B, 50 Hz: F_(19,80)_=1.2822, P=0.2188; 1 Hz: F_(19,80)_=0.3226, P=0.9964; n=5). Furthermore, there was no significant change in the latencies of ECAP and S1 across stimulus events for either 1 Hz or 50 Hz stimulation (Fig. 8C, 8D, F_(19,80)_<1, P>0.05, n=5). Therefore, S1 ESAPs amplitude are distinctly sensitive to SCS frequencies up to 50 Hz in contrast to ECAPs, suggesting a slower adapting response underlying S1 ESAPs. No visual motor responses were observed at the thresholds analyzed.

**Figure 7:**
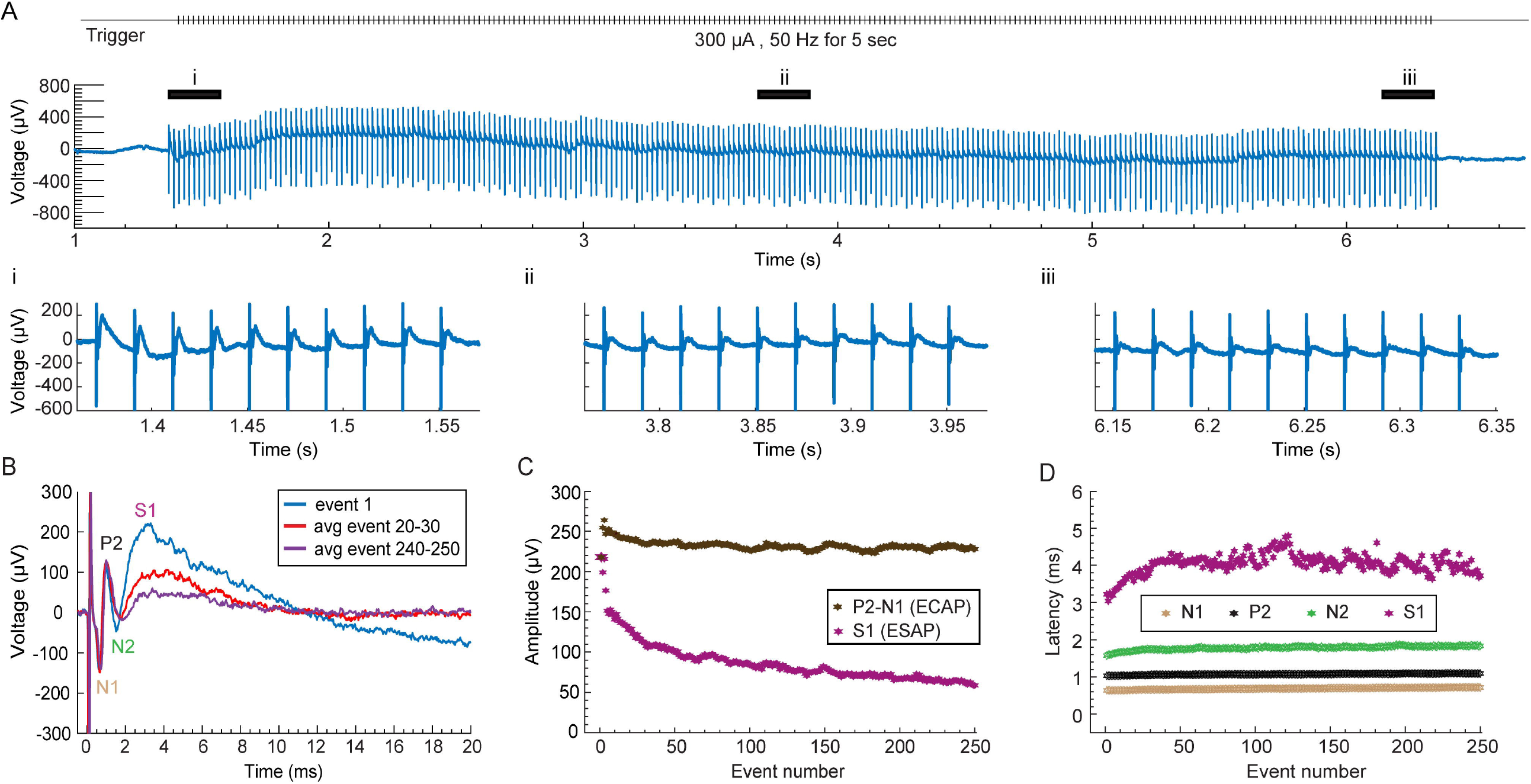
Tonic stimulation (50 Hz) evoked ECAP and ESAP (S1). **(A)** Evoked responses from the tonic stimulation of epidural spinal cord at 300 µA (40 µs pulse width, 250 stimulation events) Trigger is shown at the top. The black lines at top correspond to early (i), middle (ii), and late (iii) events shown at the bottom. The ESAP (S1) is distinctly recognizable in (i), attenuates in (ii) and (iii). **(B)** Representative traces of the evoked response comprising of ECAP and ESAP (S1) from event 1 (blue), averaged responses from event 20 to event 30 (red), and those from event 240 to 250 (purple) show a gradual decrease in the ESAP (S1) amplitude with increasing events. **(C)** Comparison of the ESAP (S1) amplitude (pink) between events. Moving average of four events is computed to smoothen the data for the amplitude of ESAP (S1) from event 7 and the amplitude of ECAP (P2-N1, brown) from event 4 onwards. Rest of the data are represented in absolute values. The ESAP (S1) amplitude drops while the ECAP amplitude stays consistent except for the first three events. **(D)** Latencies for N1, P2, N2, and S1, plotted as a function of event. Moving average of four events is computed to smoothen the data for the latencies of ESAP (S1, pink) and ECAP (N1, brown; P2, black, N2, green) from event 4 onwards. *Figure Contributions:* MS performed the experiments and analyzed the data.

**Figure 8:**
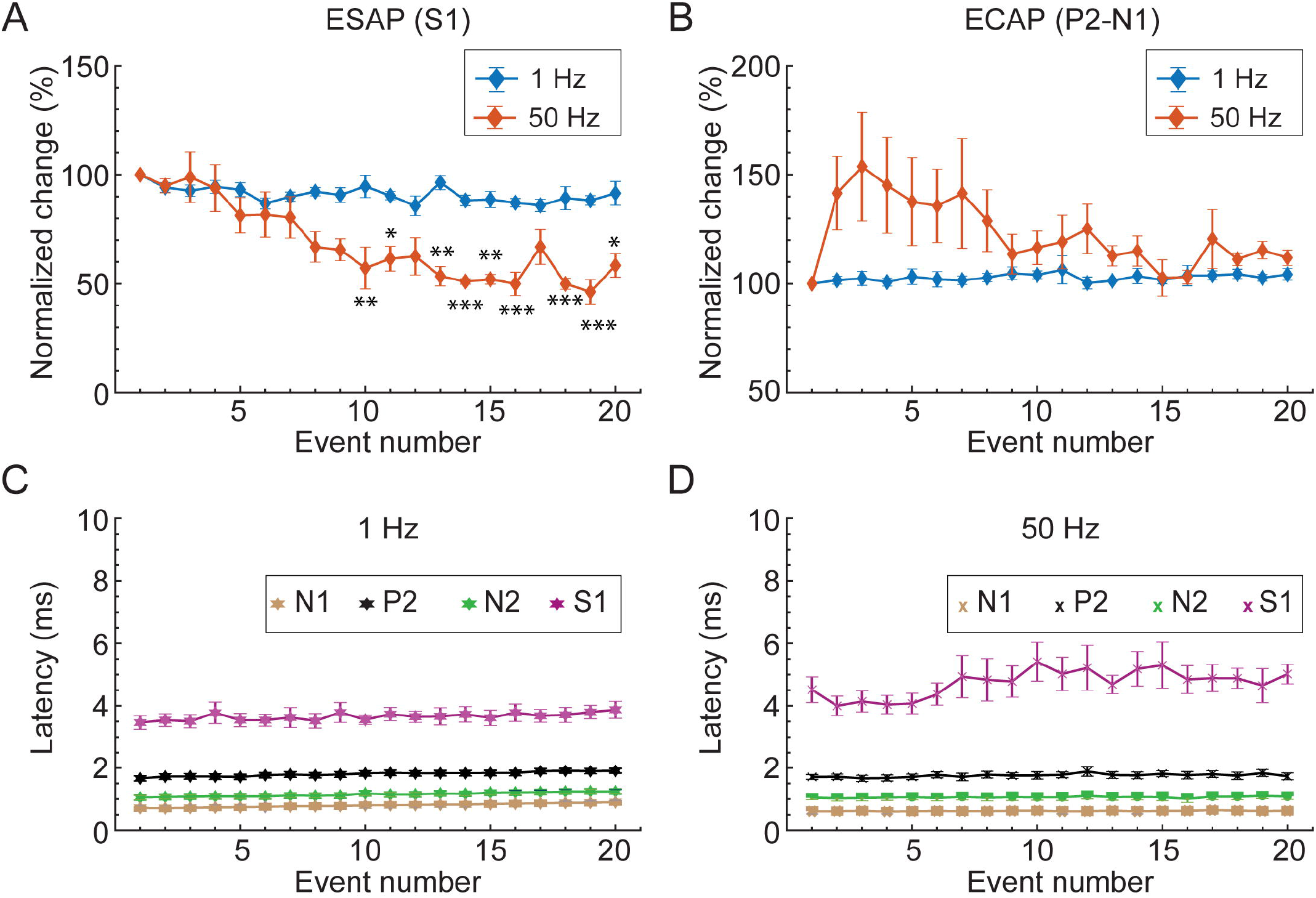
Sensitivity of ESAP (S1) and ECAP to 1 Hz vs. Tonic (at 50 Hz) SCS. **(A)** Comparison of the evoked ESAP (S1) responses from 1 Hz stimulation events (blue, Fig. 1A) vs 50 Hz SCS events (orange). Evoked ESAP responses at different stimulation events (1-20) are reported as the percentage of the ESAP value at the first event. **(B)** Comparison of the evoked ECAP responses from 1 Hz stimulation (blue) vs 50 Hz stimulation (orange). Evoked ECAP values at different stimulation events (1-20) are reported as the percentage of the ECAP value at the first event. **(C)** Latencies for evoked ECAP and ESAP: N1 (brown), P2 (black), N2 (green) and S1 (pink) for 1 Hz stimulation (n=5). **(D)** Latencies for evoked ECAP and ESAP: N1 (brown), P2 (black), N2 (green) and S1 (pink) for 50 Hz stimulation. Data represented as mean ± SEM (n=5). Statistical power using Bonferroni *post hoc* test comparing percent change in amplitudes at event 1 with those at rest of the events assumed at *P<0.05, **P<0.01, ***P<0.001. *Figure Contributions:* MS performed the experiments and analyzed the data.

## Discussion

### Features and Detection of S1 ESAPs

We show that SCS can evoke an epidural electrophysiological response termed “ESAPs” reflecting spinal synaptic currents, that are distinct (in dose response, and etiology) to well-characterized ECAPs from dorsal column axons. The observations of the S1 wave supporting this are:

a. Delayed onset and time course compared to ECAPs and N2 (Fig. 2, Fig. 5);
b. Segmental sensitivity and specificity (Fig. 2);
c. Amplitude changes and possible inversion in signal polarity across segments, in contrast to relatively fixed waveform ECAPs (Fig. 2);
d. Monotonic sensitivity to stimulation intensity, but with distinct dose response from ECAPs (Fig. 2, Extended Data Fig. 1-1);
e. Sensitivity to stimuli polarity (Fig. 2), but no direct correlation in terms of response polarity reversal, ruling against a stimulation artifact;
f. Threshold below EMG threshold (Fig. 5 B1, Fig. 6 A1), peaking before EMG initiation (Fig. 5 B2, Fig. 6 A2) and occurring independently of EMG (e.g., absence during ‘late-EMG’, Fig. 5 A1) – which along with ESAP spatiotemporal features rules out a myogenic origin;
g. Suppression by glutamatergic-synaptic antagonist (Fig. 3, Fig. 4), without concurrent block of ECAPs;
h. Diminishment (but still detectable) with frequency increases from 1 Hz to 50 Hz, in contrast to relative stability of ECAPs (Fig. 8).
i. Intraspinal correlate consistent with grey-matter origin, that is also suppressed to by a glutamatergic-synaptic antagonist (Fig. 4B).
j. Epidural electrodes measure responses to cortical stimulation, confirming detectability of spinal synaptic activity (Fig. 4C)

The ESAPs spatial distribution remains to be fully characterized as this is key to understanding the etiology of S-waves and their relevance to pain and sensory processing. Since the detection of the S1-wave is spatially constrained and dependent on the stimulation site, we suspect ESAPs reflect local connectivity (circuits), in contrast to the broadly detectable, propagating ECAPs that represent activity along dorsal column axons of passage. Thus, the absence of observable ESAPs in prior studies (Parker et al., 2013) despite visible ECAPs can be explained by variation in electrode placement. In addition, the S1-wave latency and duration approximates or exceeds the conventional SCS rates (inter-pulse intervals corresponding to 50-100 Hz frequency), that may explain why ESAPs were previously missed; along with the routine explanation that reported ECAP responses are analyzed in a limited (∼3 ms) post-stimulation pulse time window (Parker et al., 2012)(e.g. Fig. 2 and 3 insets). We further distinguish between ESAPs and multi-phasic ECAPs, as reported previously (Chakravarthy et al., 2020) and observed in our series (not shown), which reflect multiple dorsal column axon volleys, and so are distinct from S1-waves.

In contemporary SCS, limited reports of ‘late response’ spinal potentials suggested their unreliability and/or association with undesired side-effects (Falowski et al., 2022; Parker et al., 2012). We show SCS ESAPs are reliably evoked provided electrodes are positioned at specific sites, and are distinct from EMG. We also distinguish two types of SCS evoked spinal potentials labeled ‘late response’ initiating at either ∼10 ms (Parker et al., 2012) or ∼4 ms (Falowski et al., 2022). In the latter case, it was shown EMGs initiate after the ‘late response’, ruling out a myogenic origin. Muscle responses from SCS are thought to reflect activation of smaller diameter dorsal root fibers that then produce muscular activity through activation of motor neurons via reflex pathways in the ventral horn (Shils and Arle, 2012). The observation that some SCS conditions (e.g., lateral electrode placement) produce both slow spinal potentials and undesired side effects (EMG, discomfort) is therefore not surprising - and should not be confused with SCS conditions optimized for ESAPs (e.g., midline stimulation electrode at specific distance from recording electrode).

Background supporting the hypothesis that ESAPs reflect local circuit processing are wide-ranging experiments on spinal evoked (cord dorsum) potentials (Maruyama et al., 1982; Ondrejcák et al., 2005; Shimoji et al., 1982; Tomita et al., 1996; Yates et al., 1982); a ∼2 ms “triphasic spike” (attributed to axonal conduction, e.g., ECAPs), ∼5 ms “negative wave” (attributed to synaptic afferents / interneuron activity) and then a tens of ms “positive wave” (attributed to synaptic “primary afferent depolarization (PAD)” or presynaptic inhibition) have been reported. That the properties of the ESAP are sensitive to the frequency of applied SCS—with higher frequencies (i.e., 50 Hz) producing adaptation of ESAP but not ECAP—corroborates either an adapting excitatory synaptic mechanism or a result of PAD, i.e., presynaptic inhibition of excitatory afferents, as the underlying mechanism of generation. Indeed, the discovery of SCS was motivated by slow spinal responses (“after-discharges”) in the manifestation and control of pain (Shealy et al., 1970), typically recorded in the form of diffuse field potentials (Shealy, 1966). We caution against inconsistent labeling of evoked spinal responses (e.g., indexing positive and negative polarity waves) and hence adopted ‘S-wave’ here, reflecting synaptic origin (and indeed reversible polarity).

### Origin and Significance of S1 ESAPs

The occurrence of ESAPs above the thresholds of ECAPs and subsequently in time (Fig. 2), the intra-spinal grey matter sources correlated with ESAPs (Fig. 4), and the sensitivity of ESAPs to the AMPAR antagonist CNQX (Fig. 3), support the hypothesis that ESAPs reflect the synaptic excitatory inputs, and further suggest that ESAPs reflect the excitatory inputs from large diameter sensory afferents (e.g., dorsal column fibers). We present several hypotheses regarding the potential origins of ESAPs and how they may relate to the mechanisms of action of SCS or, more broadly, measure the “state” of dorsal horn activity.

ESAPs reflect rapid synaptic activation following pulses of SCS, consistent with activity from superficial dorsal horn neurons. Features of the S1 ESAP are consistent with intracellularly recorded monosynaptic responses of dorsal horn neurons to dorsal column stimulation from in vitro models (Baba et al., 1994). Neurons that receive inputs from dorsal column collaterals of large sensory fibers (e.g. Aβ afferents) include excitatory and inhibitory interneurons located in Laminae III/IV of the dorsal horn and more superficial elements (Abraira and Ginty, 2013), whose interactions and connectivity are critical to sensory and nociceptive processing (Braz et al., 2014; Peirs et al., 2015). Neurons with dendrites close and articulated toward the epidural recording electrodes could be sources for the ESAP, consistent with the intra-spinal S1-wave recording (Fig. 4B). The correspondence between signals evoked by both dorsal column and cortical stimulation (Fig. 4C), at a minimum confirm the detectability of synaptic activity originating in dorsal horn circuits and may further suggest a convergent neuronal source possessing glutamatergic synapses from dorsal column collaterals and descending cortical projections (Abraira et al., 2016).

Under the conditions tested, the ESAP (but not ECAP) exhibited gradual adaptation following 50 Hz stimulation (Fig. 7). Habituation is consistent with (though not exclusive to) synaptic processes (Sdrulla et al., 2015; Shimoji et al., 1982) and is also shown for superficial dorsal horn interneuron responses to 50 Hz SCS (Fan et al., 2022).

Alternatively, the ESAP signal may reflect activation of descending fibers (e.g., via current spread to the dorsolateral funiculus) that synapse onto the dorsal horn and, more broadly, descending modulation (distinct from antidromic segmental effects) by SCS (Saadé et al., 2015). This conjecture is consistent with the above-ECAP threshold intensities to evoke ESAPs, correspondence to cortical evoked responses, and suppression by CNQX.

All these hypotheses indicate ESAPs represent an epidurally measurable but heretofore uncharacterized indicator of dorsal horn (sensory or nociceptive) processing. Notwithstanding the motivation for further studies regarding the neural origins of the ESAP signal and translational applicability, our SCS experimental series in the context of decades of background studies on spinal electrophysiology show ESAPs may be rich sources of information on spinal cord state.

As well, ESAPs may have broader implications beyond a marker of pain modulation. Intra-spinal synaptic activity, evoked by either spinal stimulation or cortical stimulation (corticospinal tract; CST), can be detected by epidural electrodes. Where cortical or spinal electrical stimulation is used for spinal plasticity-induction (as the case for neuro-rehabilitation following injury), ESAPs can be used to provide an indicator of both acute synaptic modulation (e.g., Spike Timing Dependent Plasticity, STDP) and lasting changes in spinal motor circuits.

### Limitations

The translational applicability of ESAPs to clinical therapies requires further studies. For example, studies need to be performed to assess whether ESAPs are directly correlated with therapy and/or disease-state, as this association will elucidate their applications to neuromodulation.

Experiments were performed on anesthetized but otherwise healthy animals, so conclusions regarding ESAPs cannot be simply extended to chronic and/or neuropathic pain states. However, this limitation does not detract from the possibility that ESAPs reflect network activity in the dorsal horn gray matter and motivate future work with injury and disease models. Absolute currents thresholds will also vary across conditions (animal model, anesthesia, stimulation frequency / waveform / montage, surgical procedures etc.). Despite the absolute variability, ECAP generally had lower thresholds than ESAP (Fig. 5) and EMG thresholds are approximately ∼2x above even S1, and our reported EMG thresholds being 3x above ECAPs are consistent (or conservative) compared to prior studies (Cedeño et al., 2022; Dietz et al., 2022).

CNQX is pharmacologically selective to AMPA receptors, but cannot distinguish specific cell or fiber input types, however intrathecal CNQX delivery supports segmental rather than systemic actions, as do intraspinal recordings of the dorsal horn. It is possible that unrecorded muscle EMG contaminated the signal, but this would be inconsistent with the nature of S1 responses (e.g. inversion of polarity across segmented and within the spinal cord) and another rodent study using a similar stimulation and recording geometry confirm epidural recordings were unaffected by (local) EMG responses (Cedeño et al., 2022). We characterized one ESAP, the S1 wave, and additional ESAP signals should be considered, that may involve distinct neurotransmitters or polysynaptic connections. Indeed, our demonstration that ESAPs can be detected during SCS motivate and methodologically inform further mechanistic and application studies.

## Figure legends

**Extended Data Figure 1-1: Dose-response curve for ESAP (S1) and ECAP**. Monotonic increase in the amplitudes of ECAP (A) and ESAP (S1) (B) with increasing stimulation intensity across eight animals. (C, D) Both ECAP and ESAP (S1), when normalized to the maximum evoked response, exhibit a monotonic progression with increasing stimulation intensity. (E) The evoked ESAP (S1), when normalized to corresponding evoked ECAP, show a non-monotonic progression with increasing current. Data represented as mean of 20 trials from 8 animals (color coded).

*Figure Contributions:* MS and VB performed the experiments; MS analyzed the data.

