## Supplementary figures and images for "Novel Evoked Synaptic Activity Potentials (ESAPs) elicited by Spinal Cord Stimulation"

### Extended Data Figure 1-1

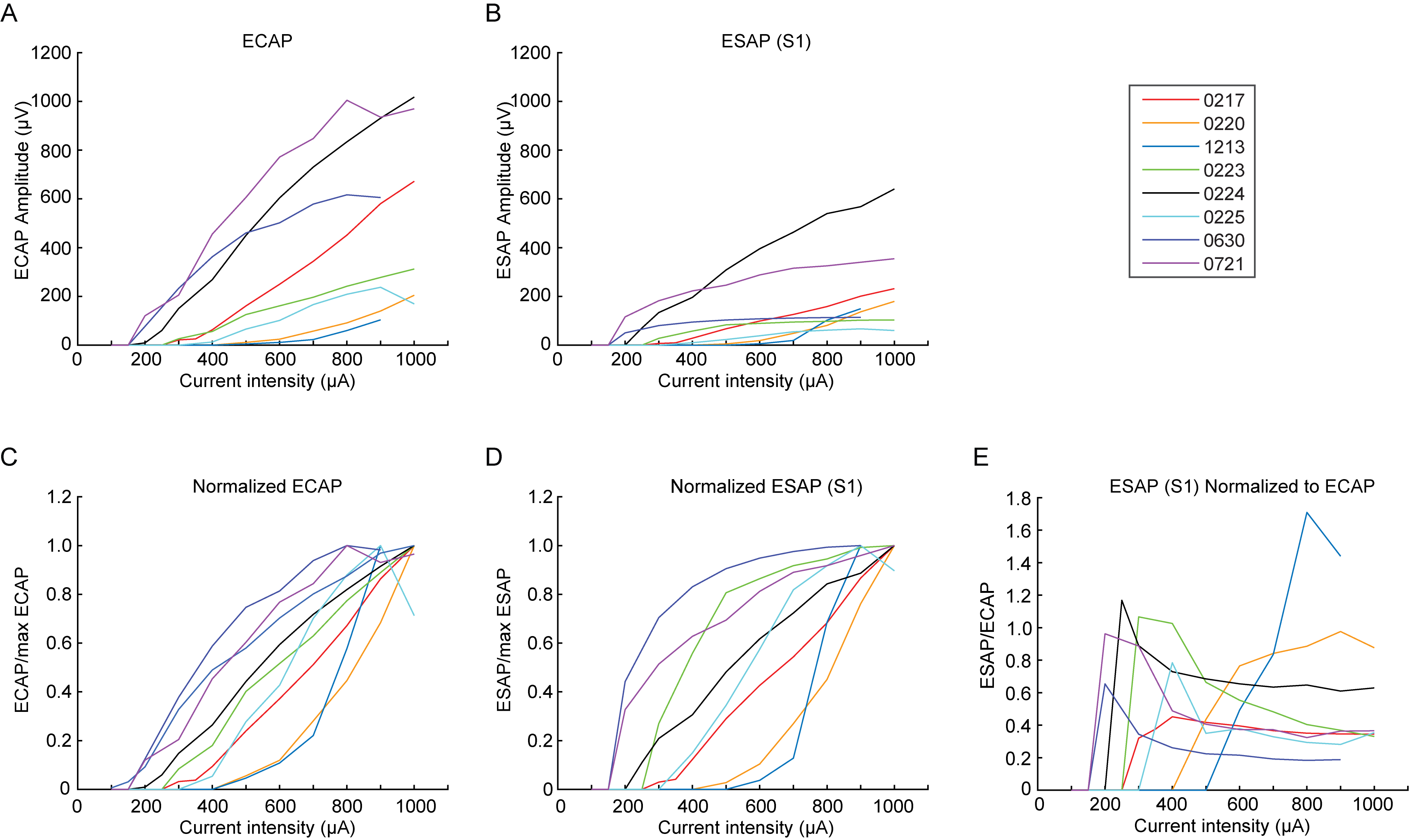
