## Supplemental Table 1 for "Novel Evoked Synaptic Activity Potentials (ESAPs) elicited by Spinal Cord Stimulation"

| Figure | Measurement | Comparison | N | Statistical test | Results |
|---|---|---|---|---|---|
| | S1 (ESAP) amplitude | Control vs CNQX | 14 | Wilcoxon signed-rank test | P=0.0001 |
| | ECAP amplitude | Control vs CNQX | 14 | Wilcoxon signed-rank test | P=0.0562 |
| | Spinal stimulation evoked S1 (ESAP) latency | Intraspinal vs epidural response | 10 | Paired sample t-test | $t_{(9)}$ = 1.35, P=0.38 |
| | Motor cortical stimulation evoked S1 (ESAP) latency | Intraspinal vs epidural response | 10 | Paired sample t-test | $t_{(9)}$ = 1.73, P=0.15 |
| | Current thresholds | S1 vs EMG | 9 | Paired sample t-test | $t_{(8)}$ = 6.04, P<0.0001 |
| | Latencies | S1 vs EMG | 9 | Paired sample t-test | $t_{(8)}$ = 15.7, P<0.0001 |
| 7A | S1 (ESAP) amplitude during 50 Hz SCS | Increasing stimulation pulse count (1-20) | 5 | One-way ANOVA | $F_{(19,80)}$=6.5162, P=8.4244e-10 |
| 7A | S1 (ESAP) amplitude during 1 Hz SCS | Increasing stimulation pulse count (1-20) | 5 | One-way ANOVA | $F_{(19,80)}$=1.3284, P=0.1898 |
| 7B | ECAP amplitude during 50 Hz SCS | Increasing stimulation pulse count (1-20) | 5 | One-way ANOVA | $F_{(19,80)}$=1.2822, P=0.2188 |
| 7B | ECAP amplitude during 1 Hz SCS | Increasing stimulation pulse count (1-20) | 5 | One-way ANOVA | $F_{(19,80)}$=0.3226, P=0.9964 |
| 7D | S1 (ESAP) latency during 50 Hz SCS | Increasing stimulation pulse count (1-20) | 5 | One-way ANOVA | $F_{(19,80)}$=0.75, P=0.7579 |
| 7C | S1 (ESAP) latency during 1 Hz SCS | Increasing stimulation pulse count (1-20) | 5 | One-way ANOVA | $F_{(19,80)}$=0.23, P=0.9996 |
| 7D | ECAP (N1) latency during 50 Hz SCS | Increasing stimulation pulse count (1-20) | 5 | One-way ANOVA | $F_{(19,80)}$=0.02, P=1 |
| 7C | ECAP (N1) latency during 1 Hz SCS | Increasing stimulation pulse count (1-20) | 5 | One-way ANOVA | $F_{(19,80)}$=1.73, P=0.049 |
| 7D | ECAP (P2) latency during 50 Hz SCS | Increasing stimulation pulse count (1-20) | 5 | One-way ANOVA | $F_{(19,80)}$=0.06, P=1 |
| 7C | ECAP (P2) latency during 1 Hz SCS | Increasing stimulation pulse count (1-20) | 5 | One-way ANOVA | $F_{(19,80)}$=0.95, P=0.5225 |
